# *Xanthomonas translucens* inhibits defense responses independently of the Type III secretion system and is recognized by cereal the immune system

**DOI:** 10.64898/2026.05.22.727309

**Authors:** Diego E. Gutierrez-Castillo, Robyn Roberts

## Abstract

Bacterial leaf streak disease (BLS), caused by *Xanthomonas translucens*, is a re-emerging disease of cereals with few effective control measures. Although the disease was identified over 100 years ago, the fundamental molecular biology and mechanisms governing host-pathogen interactions, colonization, host responses, and disease development are poorly understood. To address these knowledge gaps, we studied the early stages of host-microbe interactions between cereal hosts (wheat and barley) and *X. translucens* pv. undulosa (Xtu). We found that, while the Type III Secretion System (T3SS) is essential for disease and is associated with a 12- or 150-fold increase in bacterial populations in barley and wheat, respectively, the T3SS is not sufficient for disease development. Xanthan, an Xtu-derived exopolysaccharide, strongly contributes to symptom development by suppressing the host immune response. However, in the absence of xanthan, bacterial populations in wheat are unaffected, and in barley xanthan only accounts for a 4-fold difference compared to the wildtype Xtu. Pathogen-associated molecular pattern (PAMP) inhibition bioassays that ‘prime’ the plant immune response for subsequent infections revealed that xanthan suppresses defense priming. We also found that, in the absence of xanthan, the host immune system recognizes a potentially novel, unidentified microbe-associated molecular pattern (MAMP) of proteinaceous nature present in both *X. translucens* pathovars, undulosa (Xtu) and translucens (Xtt). Together, our work reveals both conserved and distinct *X. translucens* plant immune responses and pathogen elicitors, providing key insights into the host-pathogen interaction.

## INTRODUCTION

*Xanthomonas translucens* (Xt) is a re-emerging pathogen that causes bacterial leaf streak (BLS) disease in cereals. Originally documented in 1917, the disease has generally not been problematic until the past two decades, with a resurgence of disease focused in the upper midwestern United . BLS is found in nearly all wheat-growing regions of the world, except for some parts of Western Europe, where it is thought that the environmental conditions are not conducive to disease development (Duveiller *et al*., 1997, Paul & Smith, 1989). There is no chemical control available, and disease resistance in both wheat and barley is limited (Maraite *et al*., 2007, Ledman *et al*., 2023, Tillman *et al*., 1996, Adhikari *et al*., 2012, Duveiller, 1994). Even the efficacy of typical integrated pest management strategies is unclear (Shah *et al*., 2026). Moreover, Xt is not thought to survive long in crop debris, and thus crop rotation is likely not a major strategy to control disease (Ledman et al., 2023). The two pathovars (pv.) of major concern are pv. undulosa (Xtu), which primarily infects wheat, and pv. translucens (Xtt), primarily infecting barley. Both pathovars cause widespread disease with symptoms appearing as yellow streaks that run parallel to veins which later develop into necrotic lesions (Shah *et al*., 2023). Yield losses associated with BLS are difficult to estimate, but recent studies suggest up to a 60% loss (Friskop *et al*., 2022, Friskop *et al*., 2023).

While the disease has been recognized for over a century, there is still very little known about the basic biology or the molecular mechanisms regulating plant-microbe interactions between Xt and its hosts (Sapkota *et al*., 2020, Ledman et al., 2023). It is widely assumed that Xt is seed-transmitted, but there is conflicting evidence on whether this is the case (Ledman et al., 2023, Osdaghi *et al*., 2023). Many bacterial diseases require more humid environments and warmer temperatures to proliferate, but there is no consensus on the conditions needed for BLS disease development (Ledman et al., 2023, Sapkota et al., 2020). The two Xt pathovars (Xtu and Xtt) can infect both US major cereals, wheat and barley, but Xtu is better adapted at causing disease in wheat and Xtt is better adapted for barley. It is thought that Xtu is an intercellular pathogen and Xtt is vascular (Gluck-Thaler *et al*., 2020, Pesce *et al*., 2017). However, only one gene (*cbsA*) in Xtt/Xtu has been characterized to have adapted to different lifestyles (Gluck-Thaler et al., 2020), and it is still largely unknown what pathogen or host factors are unique to Xt and restrict host specificity.

Pathogenic microbes, including bacteria, encode a diversity of virulence factors that suppress the plant immune system. Xanthan, a mucoid exopolysaccharide (EPS) encoded by the *gum* gene suite and specific to the *Xanthomonas* family species, is a key virulence factor that is essential for disease development. *gumD* is the first gene involved in xanthan synthesis in *Xanthomonas* species, producing the monomer UDP-glucose (Revin *et al*., 2023). Most, but not all, *Xanthomonas* species produce xanthan, including *X. translucens* (Hersemann *et al*., 2017, Falahi Charkhabi *et al*., 2017), which appears as a yellow, mucoid substance on bacterial colonies. Xanthan seems to contribute to epiphytic survival on plants, perhaps partially due to its tolerance for and stability under temperature, pH, enzymatic, salt, UV and shear stresses, forming a protective biofilm matrix (Poplawsky *et al*., 2000a). It aids bacterial attachment to and movement on plant surfaces, promotes colonization of its host, and shields the bacteria from environmental stresses (Poplawsky *et al*., 2000b, Dow & Daniels, 1994, Torres *et al*., 2007, Picchi *et al*., 2021, Bianco *et al*., 2016). There is also evidence that suggests that xanthan production and quorum sensing are linked, and that quorum sensing signaling molecules regulate xanthan (EPS) production and bacterial virulence (Kanugala *et al*., 2019, He & Zhang, 2008). A *Xanthomonas campestris* quorum sensing signaling molecule (diffusible signal factor, DSF) is known to be recognized by the plant immune system, causing HR, programmed cell death, ROS production, and increased transcription of *Pathogenesis-Related 1* (PR-1). In turn, *X. campestris* suppresses these plant immune responses by secreting xanthan (Kakkar *et al*., 2015). Xanthan is also known to suppress the immune system by limiting callose deposition (Yun *et al*., 2006, Bianco et al., 2016).

It is well described that other microbial virulence factors, such as effectors, can also directly interact with plant immune receptors, blocking the immune system signaling cascade (Valenzuela & Herrera-Vásquez, 2026, Yuen & Bozkurt, 2026, Li *et al*., 2026, Maruta *et al*., 2026). Some effectors are secreted through the type-III secretion system (T3SS), a needle-like apparatus consisting of six proteins that constitute the Inner Membrane ring and export apparatus: HrcR, HrcS, HrcT, HrcU, HrcQ, and HrcV. These proteins assemble into the cytoplasmic ring (HrcC) and ATPase complex (HrcN) via an inner membrane protein (HrcJ) (Liu *et al*., 2014, Büttner, 2012). The T3SS effectors play a key role in suppressing host immunity across pathosystems. For example, tomato receptors Flagellin sensing 2 (Fls2) and Flagellin sensing 3 (Fls3), which recognize flagellin peptides flg22 and flgII-28, respectively, are inhibited by the *Pseudomonas syringae* pv. tomato T3SS effector AvrPto (Xiang *et al*., 2008, Xiao *et al*., 2007, Pedley & Martin, 2003, Shan *et al*., 2008). Transcription activator-like effectors (TALEs) modulate transcription of host RNAs to either suppress plant immunity and/or increase pathogen virulence (Mormile *et al*., 2025, Zhang *et al*., 2022, Merfa *et al*., 2025, Teper *et al*., 2023). TALEs are produced only by *Xanthomonas* species, but not all, and little is known about the targets of TALEs or how they function, especially in monocots and cereals, specifically. One TALE produced by Xtu, called Tal8, is known to affect plant immunity by targeting abscisic acid (ABA) biosynthesis (Peng *et al*., 2019). ABA is an important phytohormone and signaling molecule in plant immunity and plant abiotic stress responses, so interfering with the limiting rate-step with this pathway increases susceptibility to Xtu.

Plants defend themselves from pathogens through a variety of tactics. The plant immune system is largely classified into two broad categories, though it is now understood that these two mechanisms act synergistically (Yuan *et al*., 2021, Zhang *et al*., 2020, Yu *et al*., 2024, Ngou *et al*., 2021) Effector-triggered immunity (ETI) is activated when nucleotide-binding leucine-rich repeat receptors (NLRs) recognize specific virulence factors (effectors), typically proteins secreted by the pathogen, which initiate a cascade of molecular signaling events that results in a very strong suppression of pathogen growth through the hypersensitive response (HR). Recognition of Xt in tobacco leads to an HR response that is not observed in wheat or barley and is consistent with cereal susceptibility (Duveiller et al., 1997).While ETI is very effective at strongly restricting pathogen growth (100- to 1000-fold), it is typically species and/or strain-specific and can recognize one or a few different pathogens (Jones & Dangl, 2006). Pattern-triggered immunity (PTI) is activated when plants recognize conserved pathogen-associated molecular patterns (PAMPs), such as the bacterial motor protein flagellin, cold shock proteins, lipopolysaccharides, and chitin (Stevens *et al*., 2024). PAMPs are typically highly conserved and allow the plant immune system to broadly recognize a variety of pathogens, but generally restrict pathogen growth to a lesser extent compared to ETI (10- to 100-fold). Upon recognition of a PAMP, typically by a receptor-like kinase (RLK), global changes in gene expression occur which are associated with plant immunity. A chain of immune responses is also activated, including reactive oxygen species (ROS), mitogen-activated protein kinases (MAPKs), calcium spiking, and callose deposition. Recognition of a PAMP can also ‘prime’ the plant immune system to recognize similar PAMPs in subsequent infections, providing an effective ‘PAMP protection’ response (Desmedt *et al*., 2021, Chakravarthy *et al*., 2009). Plant defense priming agents can be biological (microbial or herbivorous), environmental (abiotic stress), or chemical (small molecules/macromolecules). Defense priming, or systemic acquired resistance (SAR), has long thought to be a mechanism through which plants may be ‘vaccinated’ against an agent, providing it ‘memory’ against a pathogen or stressor (Conrath *et al*., 2006, Westman *et al*., 2019).

Plant immunity models are largely based on data from dicotyledonous model plants such as Arabidopsis and tobacco, which are useful tools in understanding plant-microbe interactions but are not always representative of what happens in crop species (Anderson *et al*., 2005, Korwin Krukowski *et al*., 2020, Inzé & Nelissen, 2022). For example, the flagellin peptide flg22, recognized by Fls2, is an elicitor of PTI in many plant species, including Arabidopsis and *Nicotiana benthamiana*. However, a shortened version of the same peptide, flg15, is recognized only in tomato (Robatzek *et al*., 2007). Additionally, tomato Fls2 has much stronger *in vitro* kinase activity compared to the Arabidopsis homolog (Roberts *et al*., 2020, Gómez-Gómez *et al*., 2001, Cao *et al*., 2013). The tomato mitogen-associated protein kinase 3 alpha (M3Kα)-interacting 1 (Mai1) protein shares 80% amino acid identity with Arabidopsis brassinosteroid kinase 1 (Bsk1), yet is functionally divergent. In plants silenced for Mai1, HR was abolished, and this immune response could not be restored by expressing Arabidopsis Bsk1 (Roberts *et al*., 2019). Leguminous plants establish nitrogen-fixing root nodules in the presence of Rhizobia bacteria, involving regulation of common defense responses such as salicylic acid, ROS, and ethylene, yet Arabidopsis is unable to establish Rhizobial interactions (Martínez-Abarca *et al*., 1998, Mithöfer, 2002, Garrido-Oter *et al*., 2018). In wheat, transgenic co-overexpression of conserved pathogenesis-related (PR) wheat genes chitinase and a β-1,3 glucanase, or single overexpression of rice thaumatin, did not increase disease resistance against fungal pathogens despite these being well described immunity-associated genes in Arabidopsis (Hu & Rijkenberg, 1998, Anand *et al*., 2003, Chen *et al*., 1999).

While there are also many examples of the gene conservation between model plants and crop species, it is likely that many pathways and regulatory elements are different or lacking in model plants due to the genomic complexities of many crops. Therefore, while Arabidopsis and other model species provide a foundational understanding of plant immunity, crops have distinct immune features that have been shaped by evolution, plant breeding, and environmental selection. Understanding how immune systems in crops diverge from models is essential to developing plant immunity and resilience. In this study, we investigated the role of PTI and PAMPs in the Xt-wheat interaction to help elucidate the molecular mechanisms behind the wheat immune system and a re-emerging bacterial pathogen.

## METHODS

### Bacterial strains

Bacterial strains *Xanthomonas translucens* pv. translucens (Xtt CO236) (PRJNA1017868) and *X. translucens* pv. undulosa (Xtu CO237) (PRJNA1017870) (Gutierrez-Castillo *et al*., 2024a) were used for most of the inoculation experiments. Homologous recombination mutants in Xtu CO237 and Xtt CO236 were made by transforming these with TOPO™ pCR™2.1 plasmids (Invitrogen) carrying a 500bp fragment of the upstream region of *gumD* and *hrcT* (Fleites *et al*., 2013). Insertion of the TOPO™ plasmids was confirmed with either 5’ or 3’ gene-specific primers and global pUC18 primers M13F or M13R (Addgene). **Supplemental Table 1** lists all primers used in this study. We also disrupted the Xtt CO236 Type III Secretion System (Xtt CO236*ΔhrcT*) and xanthan production (Xtt CO236Δ*gumD*) using the same approach as for the Xtu CO237 mutants. Although the Xtt CO236 mutants were stable *in-vitro*, inoculation in the plant resulted in a loss of the mutant phenotype (**Supplemental Figure 1**). To confirm that the plasmid was lost and that the Xtt CO236 mutant phenotypes were not a reflection of a difference in phenotype between strains, we re-isolated bacteria from leaves syringe-infiltrated with Xtt CO236*ΔhrcT* and Xtt CO236Δ*gumD* and plated on NA agar and NA agar supplemented with kanamycin and carbenicillin (the plasmid resistance markers). Bacteria isolated from the leaves grew on NA agar but not the NA agar supplemented with the antibiotics, confirming that the plasmids were unstable and lost *in planta*.

For the PAMP protection assay, rifampicin-resistant Xtt CO236 and Xtu CO237 strains were generated. Rifampicin-resistant mutants were selected as described in (Choi *et al*., 1988). Mutants that resisted the rifampicin and appeared identical in colony morphology, growth, and disease symptomology *in planta* were selected as designated as Xtt CO236R and Xtu CO237R.

### Plant growth conditions and inoculations

Wheat (var. Hatcher) and barley (var. Morex) were grown in growth chambers at 20°C in a 16h:8h light/dark cycle with 50% humidity. Two-week-old seedlings in their second leaf stage were used for all inoculation experiments. After bacterial inoculation, plants were transferred to a growth chamber with 90% humidity, set at 22°C with 16h:8h light/dark cycle until symptom development (approximately 7 days). All inoculations were performed via syringe infiltration into the leaves. Live bacterial suspensions were prepared at an optical density (OD_600_ = 0.001) for symptom development and bacterial population assays, corresponding to approximately 10^6^ CFU/mL, unless otherwise stated.

### PAMP protection assays

PAMP protection assays were generally performed as described in (Sequeira *et al*., 1972). Bacterial solutions were prepared as in (Gutierrez-Castillo et al., 2024a, Goettelmann *et al*., 2022) at an optical density (OD_600_) of 0.1, and then heat-inactivated by subjecting the bacterial solutions to a water bath at 98°C for 10 minutes. To confirm bacterial death, the heat inactivated solutions were plated onto NA and syringe-infiltrated into wheat seedling leaves. No bacterial growth on NA plates or plant symptoms were observed (**Figure 3**). The heat-inactivated bacterial solution was syringe-infiltrated into 2-week-old wheat seedlings in their second leaf stage, and the infiltration area was marked on the affected leaves. These plants were transferred to a growth chamber set up at 22□ in a 16h:8h light/dark cycle with 90% humidity. After 24 hours, live Xtu CO237R was syringe-inoculated in the same infiltrated space as the heat-killed bacterial solution at an optical density (OD_600_) of 0.001. Symptoms were recorded 7 days post inoculation (dpi) of the live strain.

To investigate the identity of the unknown elicitor, we performed a Proteinase K (PK) digestion on the heat inactivated bacterial solutions. Proteinase K Lyophilized Powder (Thermo Fisher) was prepared according to the manufacturer’s protocol by dissolving the contents in 20mM Tri-HCl (pH 8.0), 50% glycerol, 5mM CaCl_2_. The stock PK suspension was added to the inactivated bacterial solution (OD_600_: 0.1) to a final concentration of 20mg/mL. The mixture was incubated at 37□ overnight and later stopped by setting it at 95□ for 5 minutes. The PK treated bacterial solutions were inoculated 24 hours prior to infection with the live pathogen (Xtu). Symptoms were recorded 7 dpi of the live strain.

### Xanthan recovery and infiltration

For the xanthan recovery bioassays, xanthan gum was isolated from cultures of various *Xanthomonas* species following the protocol previously described by (Dai *et al*., 2019) and prepared at OD_600_ = 0.1 with some modifications. Peptone-Yeast Malt (PYM) media (5 g dextrose, 2.5 g tryptone, 1.5 g malt extract, 1.5 g yeast extract, pH = 6.2) was prepared and sterilized by autoclaving. A single Xtu CO237 colony was used to start a culture in 250 mL of PYM and incubated at 28□ for 2 days. The bacterial pellet was recovered by centrifuging the media at 4,000 x *g* for 30 minutes at 4□. The supernatant was decanted into 50mL tubes, and 2x volumes (20mL) of 100% ethanol were added to 10mL of supernatant with 1% KCl (final concentration). The mixture of ethanol, supernatant, and KCl was swirled gently and precipitated at 4,000 x *g* for 20 minutes. This step was repeated with the rest of the supernatant to recover all the xanthan from the culture and was then left to air-dry on the benchtop for 2-3 days before dissolving the final polysaccharide pellet with deionized water. The concentration of exopolysaccharides in the last fraction was measured by optical density (OD_600_), and different dilutions of this solution were used for the xanthan recovery bioassays (**Figure 2**).

The wildtype Xtu CO237, Xtt CO236, Xtu Δ*gumD*, or water as a negative control were syringe-infiltrated with into 2-week-old wheat seedlings, and the infiltration area was marked on the leaf. The exogenous xanthan (OD_600_ = 0.1) or water was subsequently syringe-infiltrated into the same leaf area at various time points post Xtu Δ*gumD* infiltration: 0, 24, 48, and 72 hpi. Water was used as a control at each timepoint (only the 72 hpi water timepoint is shown **Supplementary Figure 3**). Symptoms were observed and photographed 7 dpi of the bacterial solution.

### Bacterial population assays

Bacterial population assays were performed in leaves as described in (Gutierrez-Castillo *et al*., 2024b). Homogeneity of variance across three independent experimental repeats was assessed using Levene’s test. Statistical significance was determined by a Kruskal-Wallis’ test followed by post-hoc multiple comparisons (p < 0.05). Data processing and statistical analyses were performed in R software using the dplyr (Wickham *et al*., 2014) and agricolae (Mendiburu, 2006). The final grouped scatterplots with overlaid mean crossbars and standard deviation were generated with ggplot2 (Wickham, 2016).

### Reactive Oxygen Species (ROS) bioassays

ROS bioassays were performed largely as described in (Hind *et al*., 2016) and (Hao *et al*., 2022) using the horseradish peroxidase (HRP)/luminol system to measure oxidative stress produced by barley plants. Specifically, for **Figure 2B**, water, Xtu CO237 (wild-type), Xtu CO237Δ*gumD*, and Xtu CO237Δ*hrcT* were infiltrated in 2-week-old barley plants. At 48 hpi, leaf discs were collected with a cork-borer size 3 (7.5mm), and placed in 200 µL of water overnight (16hr) in a 96-well polystyrene white opaque plate (Thermo Scientific). The next day, water was removed and replaced with a working solution containing HRP (20µg/mL) and luminol (34µg/mL). Either water or 100nM flg22 peptide from *Pseudomonas syringae* DC3000 were added to each well. The total Relative Light Units (RLU) accumulated over 45 minutes was measured. For **Supplemental Figure 4**, barley leaf discs from 2-week old barley seedlings were collected and suspended in water overnight (∼16 hr). Similar to the previous assay, water was removed the next day and replaced with the previously described working solution. The flg22 peptide (100nM) from *P. syringae* DC3000 (*Pst*), Xtt, or Xtu, or water (control) was added, and the total ROS (RLU) accumulated over 45 minutes was measured. For **Figure 2C**, barley leaf discs from 2-week-old barley seedlings were suspended in Xtu (OD□□□ 0.01), exogenous xanthan extracted from Xtu (OD□□□ 0.01), or water. The next day, the solutions were removed and leaf discs were resuspended in the working solution. Either water or 100nM flg22 peptide (*Pst*) were added to each well and total ROS was measured over 45 minutes.

Luminescence measurements of Relative Light Units (RLU) were collected on the SpectraMax® iD3 using an integration time of 140 ms at a sample reading distance of 9.5 mm from the top of the plate for a total of 45 minutes. The raw XML files with RLU over the 45-minute assay were converted to Excel and processed using custom scripts (Gutierrez-Castillo, 2026). Data normality was assessed using the Shapiro-Wilk test. Since the data did not fit normality assumptions, a Kruskal-Wallis test was performed followed by Dunn’s post-hoc test with Benjamini-Hochberg FDR correction (*p* < 0.05). The data processing and statistics for the ROS assays were performed using SciPy (Virtanen *et al*., 2020), pandas (The pandas Development Team) and scikit-posthocs (Terpilowski, 2019) libraries in Python 3. The Seaborn package was used for plotting the final result (Waskom, 2021).

## RESULTS

### The Type III secretion system is essential but not sufficient for disease development

The type III secretion system (T3SS) is well known to be essential for virulence for many bacterial pathogens, and is an elicitor of effector-triggered immunity (ETI). To investigate the role of the T3SS in Xtu pathogenicity, we disrupted the Xtu CO237 *hrcT* gene (Xtu CO237Δ*hrcT*) and syringe-infiltrated it or the wildtype Xtu CO237 strain into wheat or barley seedlings. In wheat, at 7 days post inoculation (dpi), the wildtype Xtu CO237 displayed the typical severe disease phenotype, including water-soaked lesions and yellowing running parallel to veins. However, the Xtu CO237Δ*hrcT* mutant nearly completely lost the disease symptomology, resulting in a complete loss of water-soaking and nearly all yellowing across the infiltrated area (**Figure 1A**). In barley, the Xtu CO237Δ*hrcT* mutant completely abolished the yellowing symptoms from a wildtype Xtu CO237 infection (**Figure 1B**). These data suggest that the T3SS is necessary for disease development.

**Figure 1.**
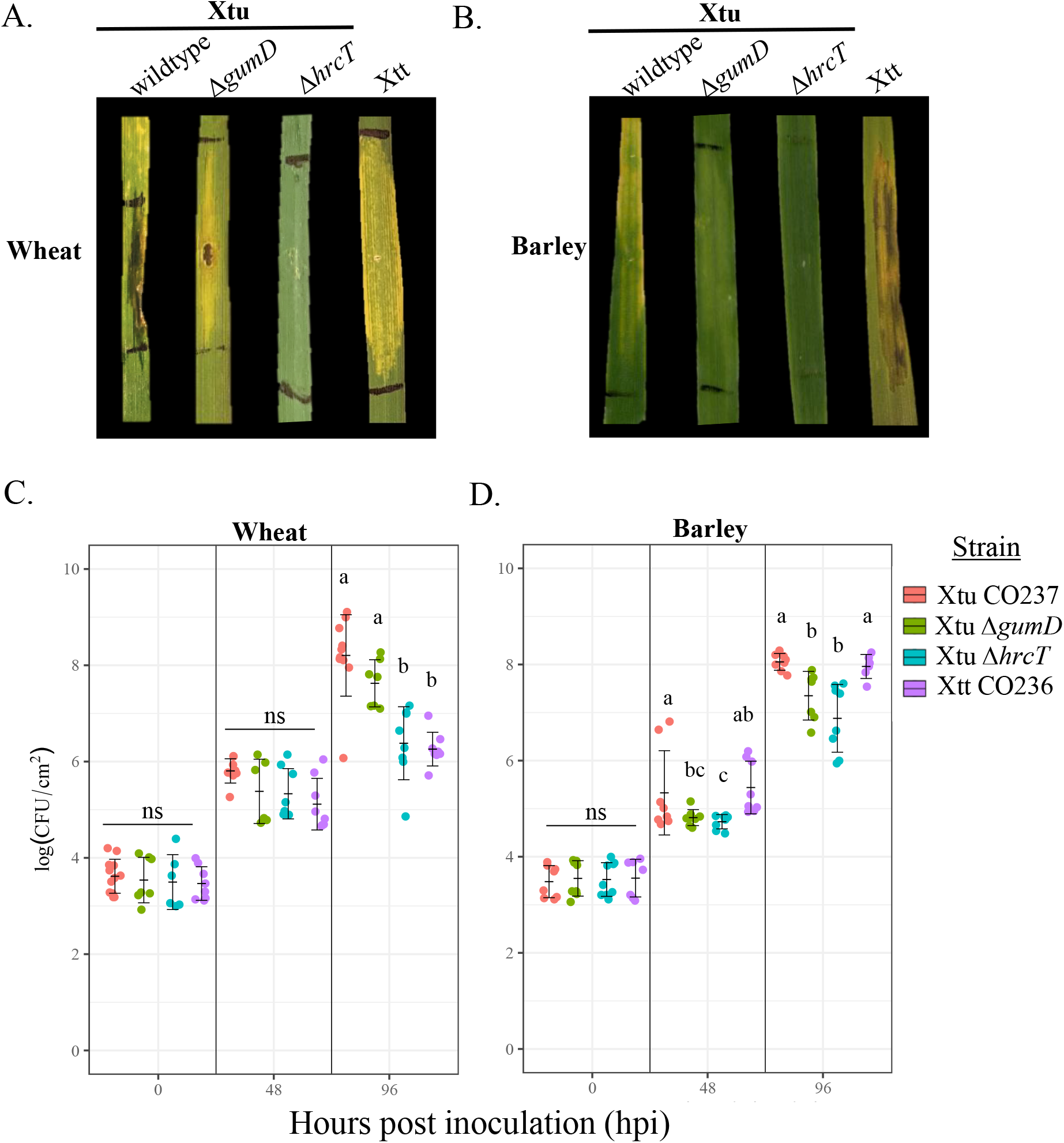
*X. translucens* pv. undulosa mutants Δ*hrcT* and Δ*gumD* have reduced virulence compared to the wild-type. A-B. Xtu CO237 (wildtype), CO237 with the type III secretion system disrupted (Δ*hrcT*), and CO237 with disrupted xanthan production (Δ*gumD*) were syringe-infiltrated into 2-week-old wheat (var. Hatcher) (**A)** or barley (var. Morex) (**B**) leaves. Photos were taken 7 days post inoculation. Experiments were repeated four times with similar results and photos from a single representative replicate are shown (n=16). **C-D**. Total bacterial populations of the wildtype Xtu CO237, Xtu CO237Δ*hrcT*, and Xtu CO237Δ*gumD*, and Xtt CO236, in wheat (var. Hatcher) (**C**) and barley (var. Morex) (**D**) leaves. Measurements were taken at 0 h, 48 h, and 96 h post syringe infiltration. Experiments were repeated three times with similar results (n=9) and compared using Levene’s test for variance (p>0.05). Significance was determined by Kruskal-Wallis test in R software (p<0.05 = significant, ns = not significant), and Tukey’s HSD was conducted to obtain different statistical groups.

We then hypothesized that xanthan, known to inhibit callose deposition, plays a role in pathogen-associated molecular pattern (PAMP)-triggered immunity (PTI). To observe the impact of xanthan production on disease development, the Xtu CO237 *gumD* gene was disrupted through homologous recombination and plasmid insertion. Mutated bacterial colonies confirmed by PCR also appeared drier and without the typical ‘mucoid’ phenotype observed with wildtype Xt, which is in alignment with a loss of xanthan production. We syringe-infiltrated the Xtu CO237Δ*gumD* mutant into wheat and barley and compared it to the wildtype Xtu CO237 and Xtu CO237Δ*hrcT*. In wheat, we observed a significant reduction of symptoms for the Xtu CO237Δ*gumD* infiltration compared to the wildtype, particularly in water soaking, but not as drastically as Xtu CO237Δ*hrcT* (**Figure 1A)**. Yellowing running parallel to the veins was still visible but did not extend beyond the infiltration area, unlike the wildtype where the yellowing continued past this region. In barley, we observed that the Xtu CO237Δ*gumD* mutant also had significantly reduced symptoms compared to the wildtype, with the yellowing symptoms much less apparent, though not as drastically as the Xtu CO237Δ*hrcT* strain **(Figure 1B)**. Knowing that the Xtt CO236 isolate causes a yellowed phenotype in wheat (but not barley), we also syringe-infiltrated Xtt CO236 into both wheat and barley to compare the yellowing phenotypes to the previously described ‘non-host’ response of Xtt on wheat. Compared to the wildtype Xtu CO237, Xtt CO236 was more similar in symptomology to the Xtu CO237Δ*gumD* mutant than the wildtype Xtu CO237. Together, these results suggest that xanthan is necessary for disease development in both wheat and barley.

### Xanthan-deficient bacterial population growth is minimally associated with symptom development

To determine if the reduction in symptoms observed for the mutants was associated with a decrease in bacterial populations, we calculated the populations in wheat and barley leaves syringe-infiltrated with each of the four strains. We measured populations (CFU/cm^2^) at 0 h, 48 h, and 96 h post syringe-infiltration in wheat (**Figure 1C)** and barley (**Figure 1D**). At 0 h and 48 h post syringe-infiltration, there was no statistically significant difference in bacterial populations between the four strains, in both wheat and barley. However, at 96 h post infiltration (hpi) in wheat, we observed that the Xtu CO237Δ*hrcT* mutant population was significantly reduced compared to the wildtype Xtu CO237. The wildtype Xtu CO237 had a mean population of 1.60 x 10^8^ CFU/cm^2^, compared to the Xtu CO237Δ*hrcT* population mean of 2.40 x 10^6^ CFU/cm^2^, reflecting a 150-fold difference. However, there was no statistically significant difference between the wildtype Xtu CO237 and Xtu CO237Δ*gumD* (population mean = 4.23x10^7^ CFU/cm^2^), nor was there a difference between Xtu CO237Δ*hrcT* and Xtt CO236 (6.25 x 10^6^ CFU/cm^2^). All four strains increased in population between 48 hpi and 96 hpi, with Xtu CO237Δ*hrcT* increased by 11-fold (p = 0.024), similar to the 14-fold difference for Xtt CO236 (1.81x10^6^ CFU/cm^2^, p = 0.003). However, the populations increased to a greater degree for the wildtype Xtu CO237 and Xtu CO237Δ*gumD* mutant with a 250-fold and 175-fold difference, respectively, between 48 hpi and 96 hpi (p<0.001).

In barley, the wildtype Xtu CO237 population increased to a similar concentration as in wheat at 96 hpi (mean population = 7.96 x 10^7^ CFU/cm^2^). However, compared to the wildtype in barley, the Xtu CO237Δ*hrcT* population was reduced by 12-fold (6.88 x 10^6^ CFU/cm^2^) (**Figure 1D)**, which is a significantly smaller difference than the 150-fold difference observed between the two strains in wheat (**Figure 1C**). The Xtu CO237Δ*gumD* mutant was also reduced slightly in barley, by 4-fold, compared to the wildtype (7.35x10^7^ CFU/cm^2^), but there was no significant difference between the two strains in wheat. Xtt CO236 in barley (8.05x10^7^ CFU/cm^2^) was not significantly different from Xtu CO237 in wheat. Between 48- and 96- hpi, bacterial populations increased for all four strains, with Xtu CO237 increasing by 330-fold (p<0.001), Xtu CO237Δ*hrcT* by 142- fold (p<0.001), Xtu CO237Δ*gumD* by 343-fold (p<0.001), and Xtt CO236 by 530-fold (p<0.001). Between 48- and 96- hpi, the four strains’ populations increased by a higher fold level in barley compared to wheat.

### Xanthan suppresses host immune responses and is necessary for leaf streak disease symptoms

To further investigate the role of xanthan in pathogen virulence, we next complemented the Xtu CO237Δ*gumD* mutant with exogenously extracted xanthan in both wheat and barley. We syringe-infiltrated the Xtu CO237Δ*gumD* mutant into wheat or barley seedlings and then syringe-infiltrated exogenous xanthan (extracted from a wildtype Xtu CO237 culture) into the same area at 0 h, 24 h, or 48 h post Xtu CO237 infiltration. We found that infiltrating xanthan 24 h post Xtu CO237Δ*gumD* infiltration complemented the mutant and restored the phenotype to levels similar to the wildtype for both wheat and barley (**Figure 2A)**. We also observed that infiltrating xanthan even later, 48 h post Xtu CO237Δ*gumD* infiltration, caused even more severe disease, beyond what we observed for the wildtype. We confirmed that xanthan or water alone did not cause a change in phenotype, nor did applying water or xanthan to the wildtype Xtu CO237 at any of these time points. Four concentrations of xanthan were tested for the complementation, and the lowest concentration that recovered the Xtu CO237Δ*gumD* phenotype to wildtype was used moving forward (OD_600_ = 0.1) **(Supplemental Figure 3)**. For controls, we used the Xtu CO237 wildtype (no treatment) to demonstrate a full recovery of the mutant phenotype. The water treatment confirms that the recovered phenotype is specific to xanthan.

**Figure 2.**
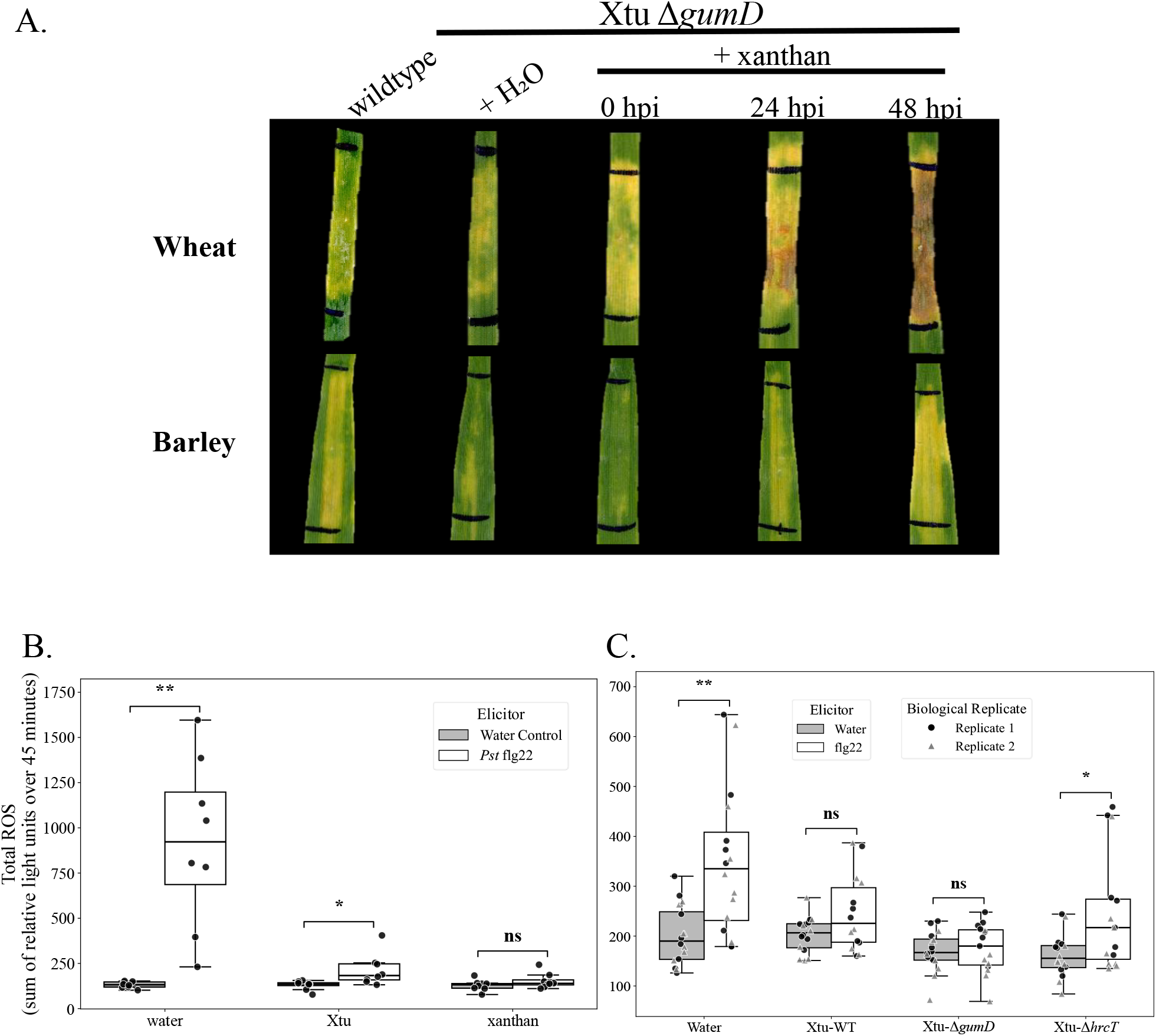
Exogenous xanthan restores the Xtu CO237Δ*gumD* mutant phenotype and suppresses the plant immune response. A. Wildtype Xtu CO237 or Xtu CO237Δ*gumD* were syringe-infiltrated into 2-week-old wheat (var. Hatcher) or barley (var. Morex) seedlings. Exogenous xanthan extracted from wildtype Xtu CO237 (OD_600_ = 0.1) or water was subsequently infiltrated into the same leaf area at various timepoints post CO237Δ*gumD* infiltration (0, 24, 48 and 72 h post infiltration (hpi) of the mutant). Photos were taken 7 days post-inoculation and are representative of three biological replicates (n=9). **B-C**. Barley leaves were either **B**. extracted for discs and floated in water, Xtu CO237 (OD□□□ _600_ = 0.01), or xanthan (extracted from Xtu CO237, OD□□□ _600_ = 0.01) for 16 h, or **C**. were infiltrated with water (control), Xtu CO237 (Xtu-WT), Xtu CO237Δ*gumD* (Xtu-Δ*gumD*), or Xtu CO237Δ *hrcT* (Xtu- Δ *hrcT)*. After 48 h, leaf discs were extracted and floated in water for 16 h. **B-C**. *P. syringae* pv. tomato DC3000 *(Pst*) flg22 peptide or water (control) was applied to elicit a ROS response. Shown is the total relative light units (RLU) summed over 45 minutes. Results are from three combined biological replicates (n=9). Control and flg22 treatments were compared for each treatment using a t-test. Significance is represented by not significant (ns) (p-value > 0.05), p-value < 0.05 (*), and p-value < 0.005 (**).

The flg22 peptide from bacterial flagellin is a known elicitor of reactive oxygen species (ROS) responses in many plants elicit a ROS burst, so we wanted to determine if flagellin from Xtu CO237 or Xtt CO236 could activate an immune response. Therefore, we tested whether flg22 from Xtu CO237, Xtt CO236, or *Pseudomonas syringae* pv. tomato (*Pst)* strain DC3000 could cause this response in wheat and barley. We were unable to detect ROS in wheat using a traditional ROS assay, so we moved forward with conducting the assays in barley. Barley responded to flg22 from *Pst* DC3000, but not Xtt or Xtu **(Supplemental Figure 2**), which is in alignment with previous studies (Stevens et al., 2024).

To investigate whether xanthan could block a plant immune response, we took leaf discs from barley and suspended them in water (negative control), Xtu CO237, or xanthan overnight. The next morning, water (negative control) or the flg22 peptide was added to each of the treated primed discs to elicit ROS. We observed no ROS burst in response to the xanthan-primed flg22 peptide (**Figure 2B**). This suggests that xanthan blocks the ROS response elicited by flg22 peptide, either directly or through defense priming.

To further investigate this response, we syringe-infiltrated barley leaves with water, the wildtype Xtu CO237, Xtu CO237Δ*gumD*, or Xtu CO237Δ*hrcT* (control) and collected leaf discs 48 h later. After floating the discs in water overnight, flg22 peptide or water (control) was added to the discs and ROS responses were measured. As expected, the discs infiltrated with water responded to flg22 but not water, indicating that neither water nor the damage caused by the syringe infiltration primed the immune response **(Figure 2C)**. The leaves infiltrated with wildtype Xtu CO237 had a reduced flg22 response, comparatively. While there was a slight observable difference between the water and flg22 treatments primed with wildtype Xtu CO237, this was not statistically significant (p=0.069). Because T3SS effectors often inhibit the plant immune response and are also typically elicitors of immunity, the absence of the T3SS results in no defense priming and no immunity inhibition. As expected, when Xtu CO237Δ*hrcT* was infiltrated, a flg22 response was induced compared to the water control (p=0.020). However, leaf discs infiltrated with Xtu CO237 Δ*gumD* did not respond to flg22, and this response was not significantly different from water (p=0.717). Together, these data suggest that xanthan alone does not activate the plant immune system, and xanthan inhibits plant immunity 24-48 h post bacterial infiltration.

### A proteinaceous factor from Xtu and Xtt primes the wheat and barley plant immune systems

To investigate whether a Xtu CO237 factor is recognized by the wheat and barley immune systems, we primed the immune system by infiltrating heat-inactivated (boiled) bacteria in wheat and barley leaves 24 hours prior to infiltrating the same area with live bacteria (**Figure 3**). We boiled Xtu CO237 and Xtt CO236 and confirmed they were inactivated by infiltrating them into wheat and barley, where we observed no disease. To induce defense system priming, we infiltrated heat-inactivated Xtu or Xtt into wheat or barley leaves, then infiltrated live Xtu CO237 inoculum into the same area either 0 h or 24 h later. In wheat, we observed that infiltrating either heat-inactivated Xtu or Xtt 24 h prior to live Xtu resulted in lessened disease symptoms compared to the wildtype infiltrated with water. In fact, live Xtu primed with heat-inactivated Xtt caused slightly less disease than heat-inactivated Xtu. In barley, we also saw fewer symptoms when inactivated Xtu or Xtt was infiltrated 24 h prior to live Xtu, and inactivated Xtu provided slightly better protection than heat-inactivated Xtt. This suggests that a bacterial factor, independent of live metabolism, is recognized by the wheat and barley immune systems.

**Figure 3.**
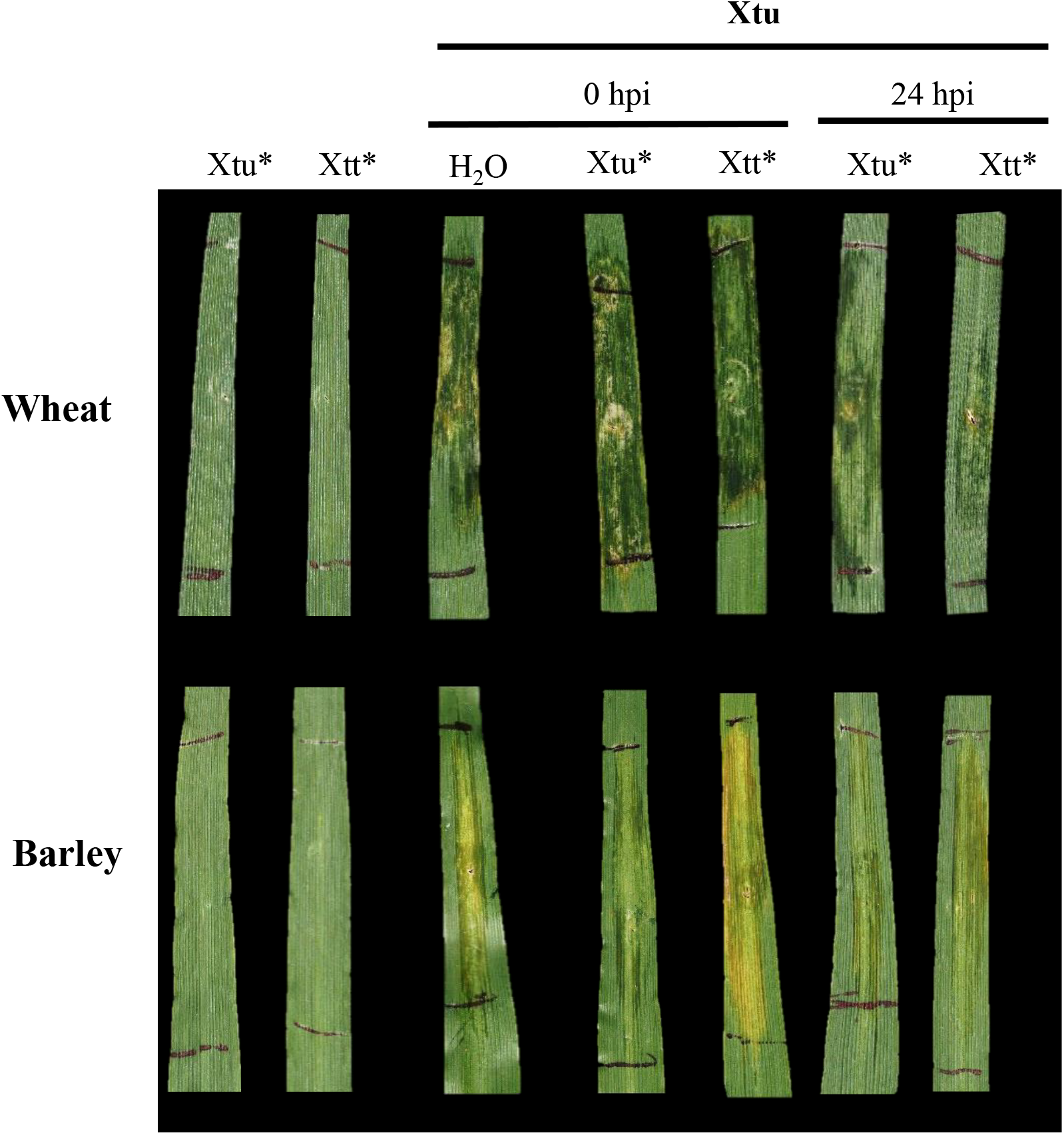
Heat-inactivated bacterial suspensions suppress *X. translucens* pv. undulosa symptoms in both wheat and barley. Xtu CO237 and Xtt CO236 were heat inactivated (*) and syringe-infiltrated into 2-week old wheat seedlings (var. Hatcher) or barley (var. Morex) leaves. Water was infiltrated as a negative control, and the Xtu* and Xtt* were infiltrated alone as a control to ensure bacteria were inactive. At 0- or 24- hours post infiltration (hpi) of Xtu* or Xtt*, live Xtu inoculum was infiltrated in the same leaf area, denoted by black lines. Photos were taken 7 days post inoculation of the live Xtu pathogen. Data shown are from a single biological replicate and are representative of a two replicates (n=8).

We hypothesized that the elicitor of the cereal defense priming response is proteinaceous in nature. To test this, we subjected the heat-inactivated bacterial solutions (Xtu and Xtt) to a protein stability test using the serine protease Proteinase K (PK). We treated the heat inactivated Xtu or Xtt solutions with or without PK to cleave peptide bonds present in the suspension, and used water as a negative control. Heat-inactivated Xtu or Xtt solutions treated or untreated with proteinase K were syringe-infiltrated into wheat or barley leaves. 48 h later, live Xtu was infiltrated in the same area and symptoms were observed after 7 days. In both wheat and barley, we observed that untreated, heat inactivated Xtu solutions infiltrated 48 h prior to live Xtu inoculations developed less severe symptoms than the Xtu wildtype strain pre-infiltrated with water (control) (**Figure 4**). However, the PK-treated, heat inactivated bacterial solution displayed more severe symptoms compared to the untreated heat-inactivated Xtu pre-infiltration, which more closely resembled the control. A similar result was observed using the PK-treated, heat inactivated Xtt extract in both wheat and barley. Compared to the controls, the leaves primed with PK-treated, inactivated Xtt had more severe symptoms than the untreated inactivated Xtu. These data indicate that the elicitor of wheat and barley defense priming is proteinaceous in nature.

**Figure 4.**
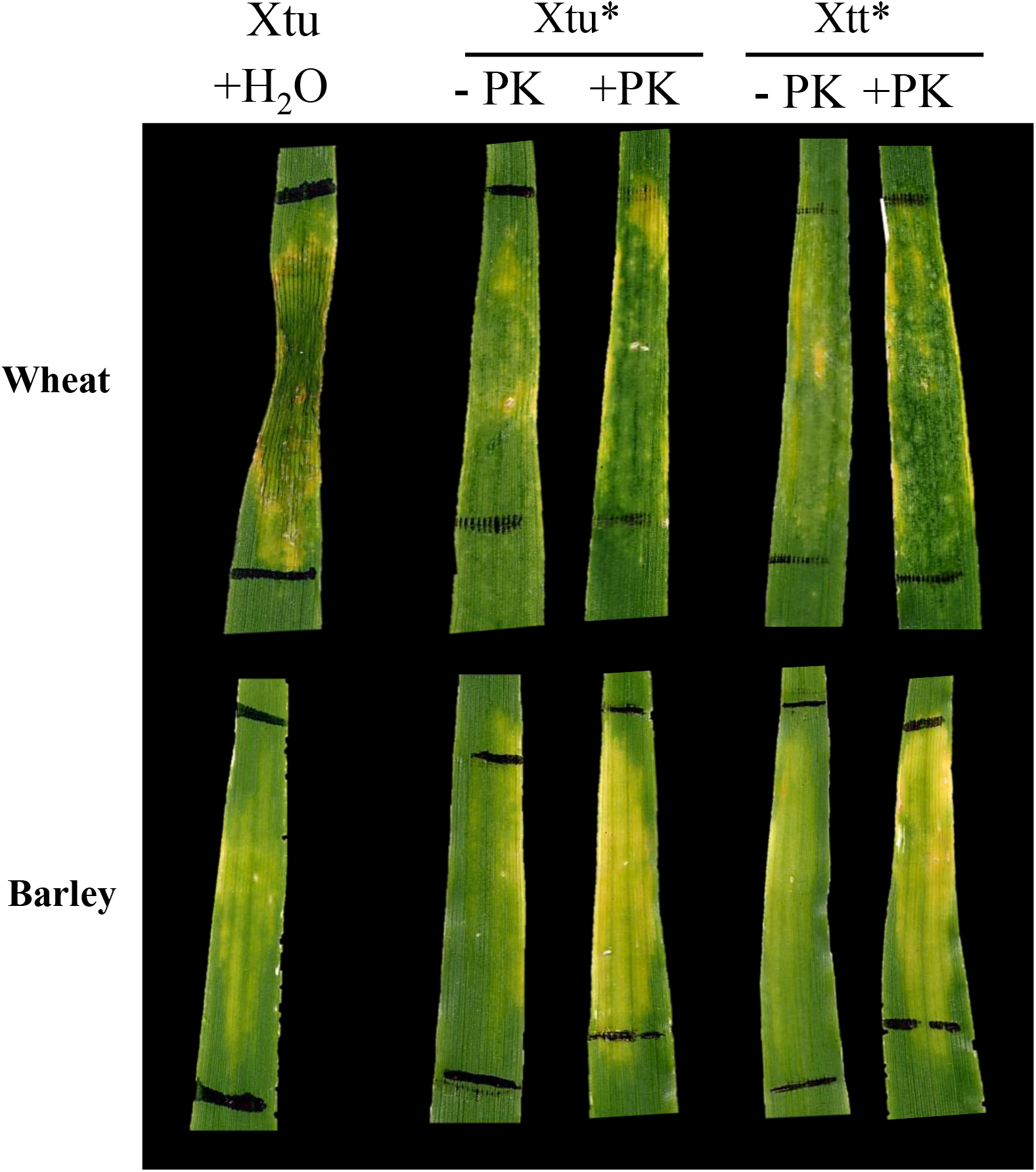
Treating heat-inactivated bacterial suspensions with proteinase K restores the Xtu symptoms. Xtu CO237 and Xtt CO236 were heat inactivated (*) and subjected to a proteinase K (PK) digestion. Non-digested Xtu* and Xtt* were included as controls. These Xtu* and Xtt* treated (+PK) or untreated (-PK) solutions were then syringe-infiltrated into wheat (var. Hatcher) or barley (var. Morex) leaves, and water was infiltrated as a negative control. At 24- hours post infiltration (hpi) of Xtu* or Xtt* +/- PK, live Xtu was infiltrated in the same leaf area, denoted by black lines. Photos were taken 7 days post inoculation of the live Xtu pathogen. Data shown are from a single experiment and are representative of a two biological replicates (n=8).

## DISCUSSION

The molecular mechanisms controlling immunity in cereals are largely unknown, as well as their interactions with pathogens including *Xanthomonas translucens*. Here, we investigated some of the requirements for *X. translucens* virulence, including the role of xanthan in disease development and the inhibition of defense activation, observed Xtu and Xtt bacterial population growth during the initial days of infection, and discovered that both wheat and barley recognize a proteinaceous factor from Xtu and Xtt that elicits an immune response. Together, these findings provide foundational knowledge of molecular wheat-bacterial interactions for a pathogen that is re-emerging with no significant disease resistance or chemical controls.

We found that the T3SS is required for disease development in both wheat and barley, suggesting that effectors secreted by the T3SS play essential roles in disease development (**Figure 1A)**. This is a widely-used strategy for many bacterial pathogens, including Xanthomonads, which use both canonical and TAL effectors to support their virulence (Phan *et al*., 2025, Kay & Bonas, 2009, van der Hoorn & Kamoun, 2008, Macho, 2016, Teper et al., 2023, Zhang et al., 2022). Similar to other Xtu and Xtt isolates, Xtu CO237 has 31 canonical effectors and 8 TAL effectors, and Xtt CO236 has 34 canonical effectors and 5 TAL effectors (Goettelmann et al., 2022, Gutierrez-Castillo et al., 2024a). Both of these strains are highly aggressive, and the loss of symptoms when the T3SS is deleted suggests that effectors are required for leaf streak disease development. However, while the T3SS was essential for maximum bacterial populations, it was not required for population growth. Xtu CO237Δ*hrcT* populations increased to similar levels in wheat and barley, and both increased exponentially over time with mean populations of 2.40 x 10^6^ CFU/cm^2^ and 6.88x10^6^ CFU/cm^2^, respectively, 96 h post infiltration (**Figure 1B**). This suggests that T3SS elements, likely effectors, may be essential for virulence, which contribute to pathogenicity but may also be recognized by the plant immune system. The 150-fold difference in wheat between the Xtu CO237Δ*hrcT* mutant and the wildtype are also in alignment with the 100-1000-fold reduction in populations observed from ETI. In barley, there was a 12-fold difference between the Xtu CO237Δ*hrcT* mutant and the wildtype, which may suggest that Xtu effector suites play a greater role in pathogen virulence in wheat over barley. Additionally, effectors may participate in immune suppression since the Xtu CO236 Δ*hrcT* mutant did not elicit immunity priming in barley ROS assays **(Figure 2B)**. Together, this may indicate that the Xtu CO237Δ*hrcT* mutant may have a significant fitness cost due to the loss of effectors but can also avoid the plant immune system, allowing populations to increase without causing disease.

Xanthan plays an important role in disease development. Xtu CO237Δ*gumD* developed less severe symptoms, particularly in water-soaking, in both wheat and barley compared to the wildtype **(Figure 1A**), and exogenously added xanthan could recover the wildtype phenotype (**Figure 2A)**. It seems to be most important around 24-48 h post infection, which may primarily be due to its role in suppressing earlier stages of the host immune system. In barley, we saw that Xtu CO237 and xanthan suppressed the flg22 ROS response **(Figure 2B**). The Xtu CO237Δ*gumD* mutant infiltrated 48 h prior to flg22 elicitation permitted defense priming and resulted in no ROS burst from flg22 **(Figure 2C**). There was a small observable difference in the wildtype ROS response between water and flg22, though it was not statistically significant. This may be because xanthan does not fully inhibit the immune response specifically linked to the ROS burst, leading to an incomplete reduction in immunity responses. It is also possible that xanthan may more strongly inhibit other immunity pathways independent of ROS. Additionally, there was a small reduction (4-fold) in bacterial populations of Xtu CO237Δ*gumD* compared to the wildtype in barley, but no statistically significant difference between the two strains in wheat (**Figure 1B**). While xanthan is important for disease development and suppressing the plant immune system in both wheat and barley, there may be some divergence in the role of xanthan in bacterial populations. Xanthan likely plays a role in quorum sensing, since quorum sensing molecules regulate xanthan production and bacterial virulence (Kanugala et al., 2019, He & Zhang, 2008). Perhaps, in barley, xanthan is more closely linked to quorum sensing and the xanthan/quorum sensing signaling increases with bacterial replication.

A boiled, heat-inactivated solution of Xtu CO237 or Xtt CO236 could both prime the wheat and barley immune system against a subsequent Xtu CO237 infection **(Figure 3**). Interestingly, Xtu primed barley more effectively than Xtt, but Xtt primed wheat better than barley. Knowing that the elicitor is proteinaceous in nature, since a proteinase K digest reduced the priming effect (**Figure 4**), there may be differences in the structure (fold), binding efficiencies, or sequence of the elicitor that are more effectively recognized by one host over the other. Investigating the differences between the Xtu and Xtt elicitors and their host receptors, and the molecular mechanisms inducing the responses, will shed light on monocot immune responses. Understanding these interactions may also lead to engineered solutions to disease control and prevention.

Xtu and Xtt typically cause the most severe disease on the host from which they were isolated, but many isolates cause lesions on both wheat and barley (Curland *et al*., 2018). In wheat, Xtt typically causes a ‘yellowing’ symptom when inoculated via syringe. While this response has been previously described as a ‘non-host’, ‘non-pathogenic,’ or resistance response, there is still little experimental data to support this. The response is unlike a typical resistance or hypersensitive response, nor does it clearly represent disease; therefore, this phenotype is yet undefined (Pesce et al., 2017, Curland *et al*., 2020, Adlung *et al*., 2016, Balint-Kurti, 2019, Dubrow *et al*., 2022, Sapkota et al., 2020). Non-host resistance typically completely restricts pathogen replication, or allows pathogens to replicate minimally before limiting it, and accounts for a 10-100-fold difference in population numbers (Gill *et al*., 2015, Schulze-Lefert & Panstruga, 2011, McLellan *et al*., 2022, Nagaraj *et al*., 2016, Coemans *et al*., 2008). Notably, when we inoculate Xtt CO236 into wheat, there is a high bacterial population present (mean = 1.81x10^6^ CFU/cm^2^ for Xtt in wheat 96 h post-infiltration) that increases exponentially but causes the ‘yellowing’ phenotype (**Figure 1)**. The phenotype closely resembles the Xtu CO237Δ*gumD* mutant, which we hypothesize is unable to block plant innate immunity due to the lack of xanthan, which may provide some support for the phenotype reflecting an undefined resistance-type response. A recent pre-print suggests that a single effector may be responsible for the ‘yellowing’ response, and they observe similar bacterial population levels for their Xtu Δ*hrcT* mutant compared to our study (Merfa et al., 2025). Future studies focusing on the cause of the yellowing phenotype will provide insights into the mechanisms behind symptom development.

## ACKNOWLEDGEMENTS

We thank Dr. Jonathan Jacobs and Dr. Veronica Roman-Reyna for providing the *gumD* and *hrcT* TOPO plasmids, and Colorado State University undergraduates Libby Swanson, Emma Barrett and Sidney Young for their technical support. This work was funded by the Colorado Wheat Administrative Committee, USDA Extension Implementation Program grant (#70006-43565), and Colorado State University start-up funds.

## AUTHOR CONTRIBUTIONS

*Robyn Roberts*: Conceptualization, Writing-Original Draft, Writing-Review and Editing, Supervision, Project Administration, Funding Acquisition, Validation, Visualization. *Diego Gutierrez*: Conceptualization, Writing-Original Draft, Writing-Review and Editing, Methodology, Formal Analysis, Data Curation, Validation, Visualization, Investigation.

