## Supplemental Figures for "*Xanthomonas translucens* inhibits defense responses independently of the Type III secretion system and is recognized by cereal the immune system"

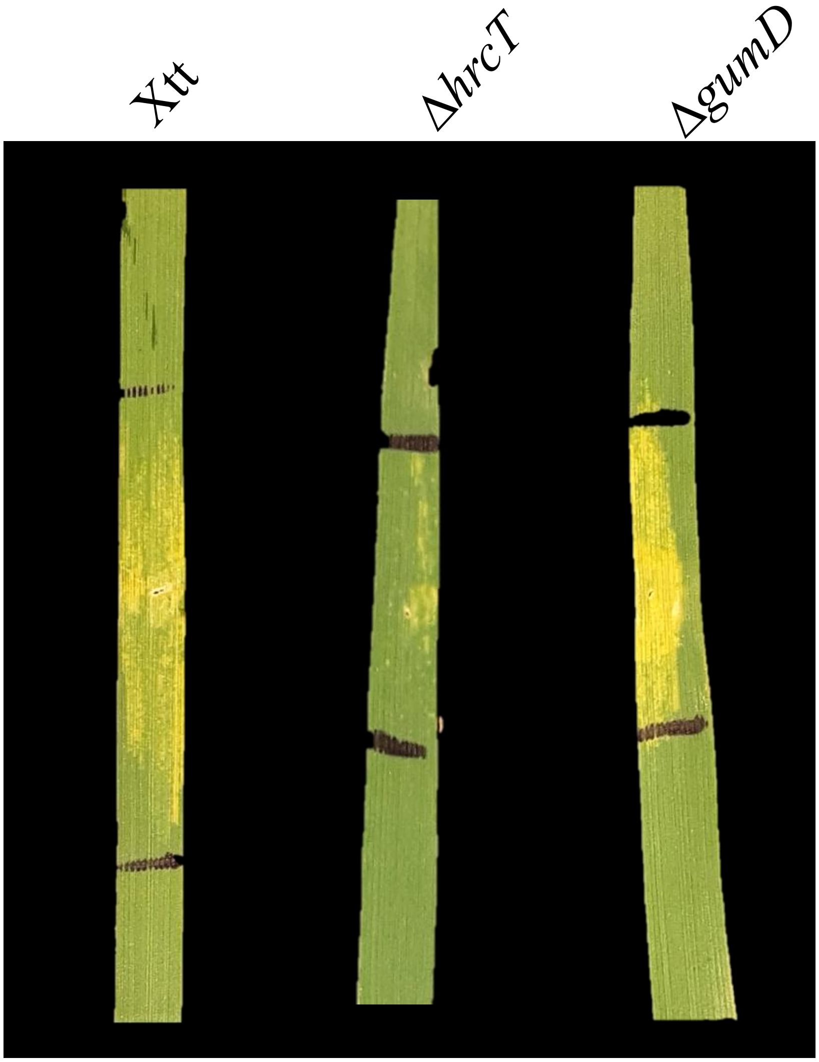


**Supplementary Figure 1. The Xtt CO236 mutants Xtt CO236Δ*hrcT* and Xtt CO236Δ*gumD* are unstable in wheat seedling leaves.** The wildtype and mutant strains were syringe-infiltrated into wheat seedling leaves. Colony PCR and an antibiotic screen confirmed that the mutants lost the mutations after infiltration. Photos are representative of a single biological replicate (n=4) and were taken 7 days post inoculation of each strain.

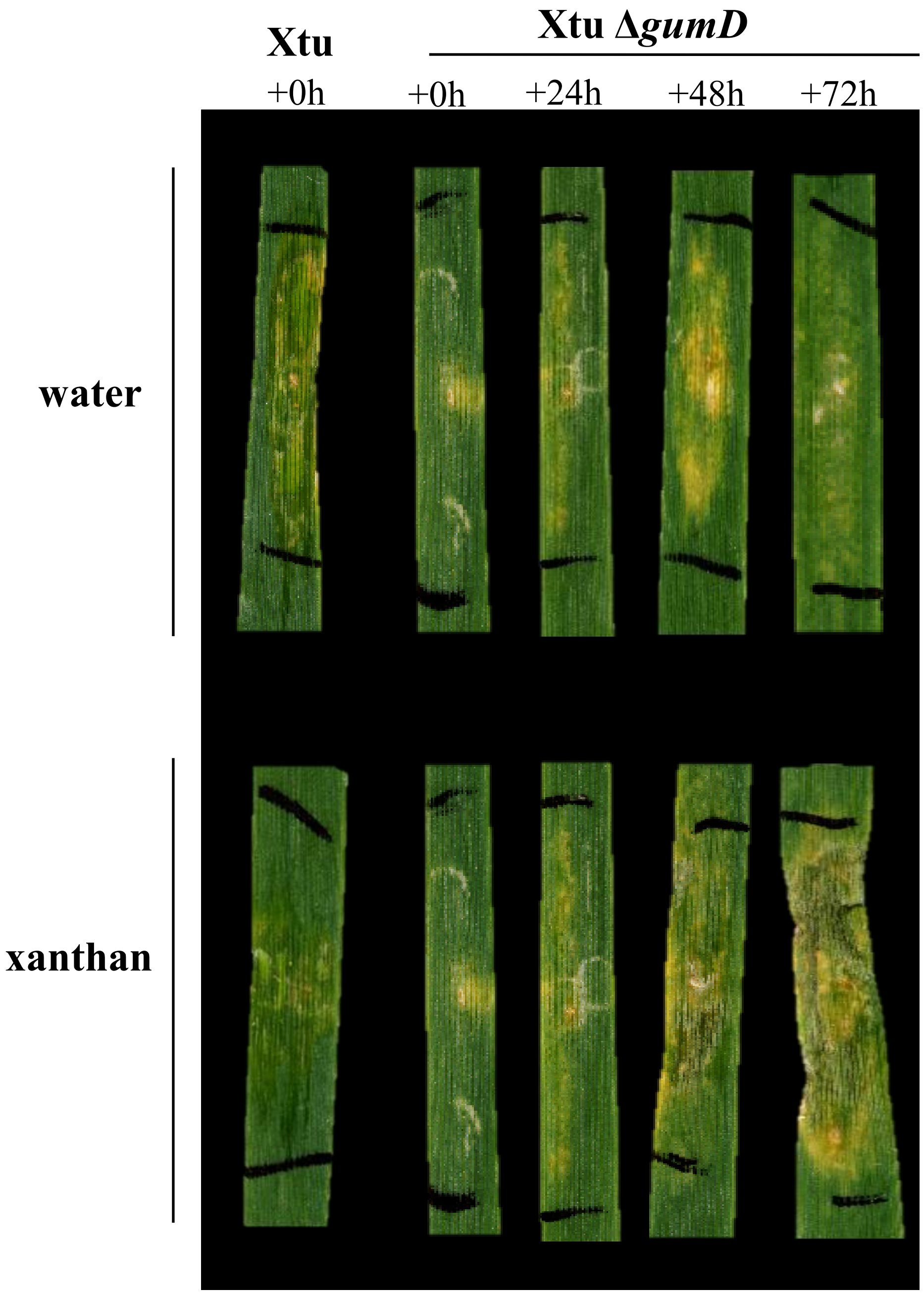


**Supplementary Figure 2. The Xtu CO237Δ*gumD* phenotype is specifically restored by exogenously-added xanthan at 24-48 h post bacterial infiltration.** Xanthan or water (control) was infiltrated 0, 24, 48 or 72 h post syringe infiltration (hpi) of Xtu CO237Δ*gumD*. The Xtu CO237 wildtype strain was syringe infiltrated and included as a positive control. Photos are from a single biological replicate and were taken 7 days post inoculation of the wild-type and Xtu CO237 Δ*gumD* (n=4).

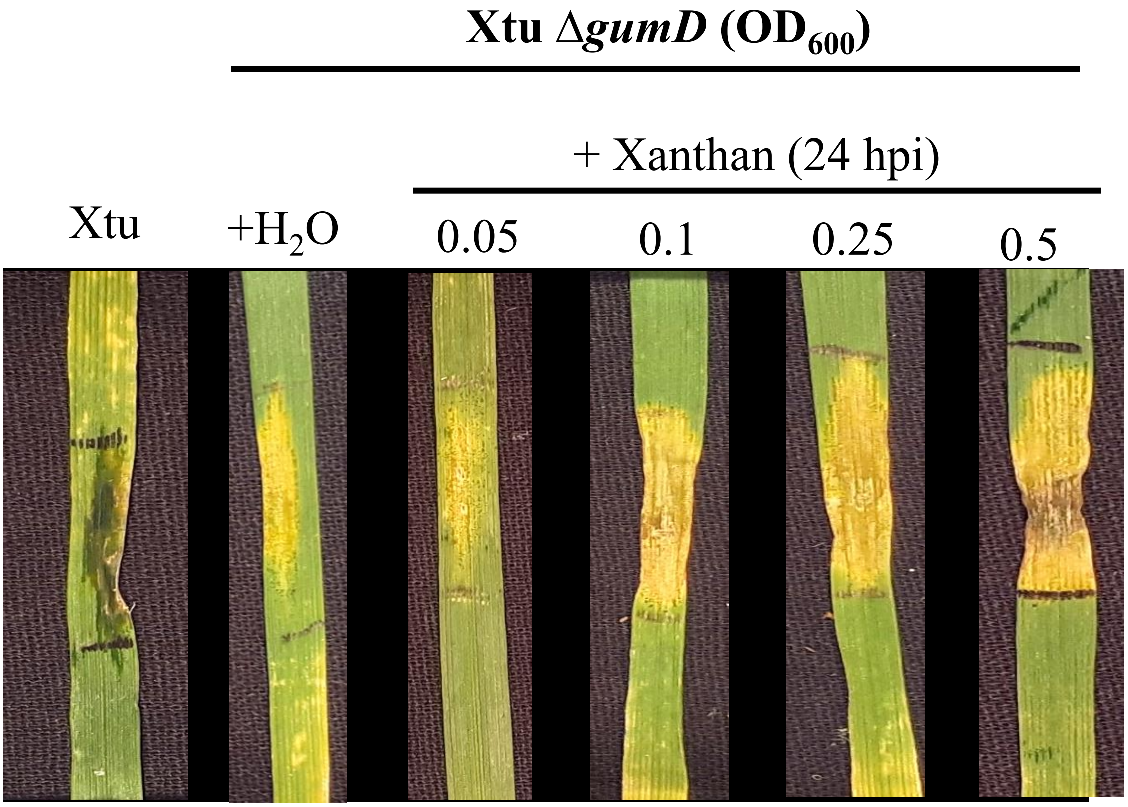


**Supplementary Figure 3. Xanthan ($OD_{600}$ = 0.1) complemented the Xtu CO237Δ*gumD* mutant phenotype**. Wheat seedling leaves were syringe-infiltrated with Xtu CO237Δ*gumD* or Xtu CO237 wildtype (control). At 24 hours post infiltration (hpi), xanthan at $OD_{600}$ = 0.05, 0.1, 0.25, or 0.5 was infiltrated into the same leaf area as the wildtype or mutant bacteria strains, denoted with black lines. Photos were taken at 7 days post-inoculation and are representative of four plants for each treatment (n=4).

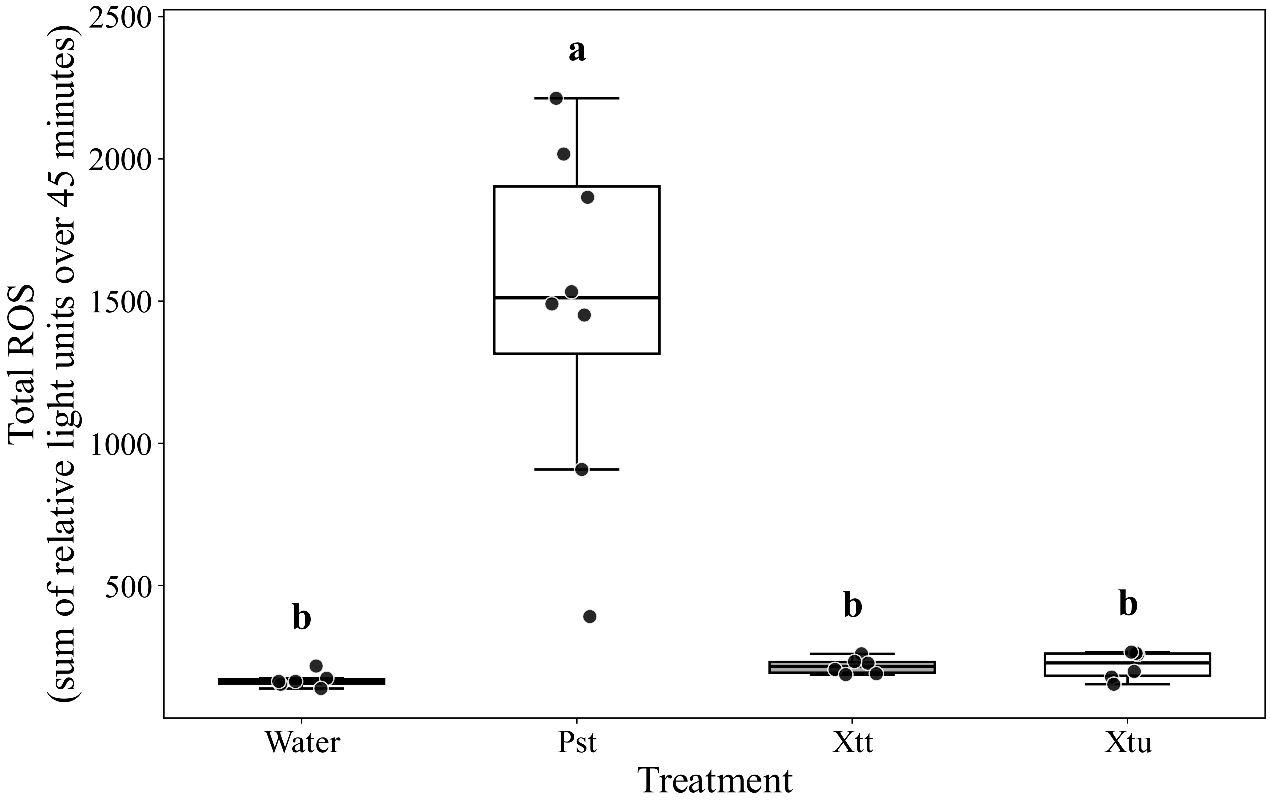


**Supplementary Figure 4. Barley does not activate a ROS burst in response to the Xtu or Xtt flagellin peptide flg22.** Leaf discs from barley seedlings were extracted and floated in water for 16 h before eliciting an immune ROS burst using Xtu CO237, Xtt CO236, or *P. syringae* pv. tomato *(Pst)* flg22 peptide, or water as a control. Shown is the total relative light units (RLU) summed over 45 minutes. Results are from three combined biological replicates (n=9). Data was tested for normality using a Levene's test ($p>0.05$), followed by and an ANOVA ($p<0.05$) and Tukey's HSD to obtain different statistical groups.
